# CAIX-targeted α therapy directed against hypoxic tumor cells in combination with immune checkpoint inhibitors in a syngeneic mouse tumor model

**DOI:** 10.64898/2026.01.06.697674

**Authors:** Sylvia T.M. Wenker, Mark W. Konijnenberg, Daphne Lobeek, Giulia Tamborino, Janneke D.M. Molkenboer-Kuenen, Gerben M. Franssen, Daan F. Boreel, Simone C. Kleinendorst, Hans Peters, Johan Bussink, Sanne A.M. van Lith, Sandra Heskamp

**Author notes:** Corresponding author: Sylvia Wenker, Geert Grooteplein Zuid 10.

## Abstract

Tumor hypoxia is a major factor in therapy resistance. A potential strategy to treat hypoxic tumors is targeted α therapy (TAT), since α particles can cause complex DNA damage independent of oxygen levels. Here, we investigate the potential of TAT as monotherapy and in combination with immune checkpoint inhibitors (ICI) to treat hypoxic tumors.

**Methods:** Monoclonal anti-CAIX antibody DOTA-MSC3 was labeled with indium-111 (^111^In) or actinium-225 (^225^Ac), and binding to CAIX-expressing hypoxic tumor cells was determined *in vitro*. Subsequently, the *in vivo* biodistribution and dosimetry of radiolabeled DOTA-MSC3 was assessed in B16F10-OVA tumor-bearing mice, and its spatial distribution in the tumor (autoradiography) was correlated to CAIX expression measured by immunofluorescence. Finally, tumor growth and survival were determined upon treatment with [^225^Ac]Ac-DOTA-MSC3 with and without ICI.

**Results:** ^111^In- and ^225^Ac-labeled DOTA-MSC3 bound specifically to CAIX-expressing hypoxic tumor cells. *In vivo*, uptake of both radiopharmaceuticals in B16F10-OVA tumors was spatially correlated with CAIX-positive hypoxic tumor regions. [^225^Ac]Ac-DOTA-MSC3 significantly prolonged survival of mice compared with PBS control (p=0.0032). Furthermore, the combination of [^225^Ac]Ac-DOTA-MSC3 and ICI significantly delayed tumor growth and prolonged survival compared with PBS control (p=0.0022 and p=0.0019, respectively).

**Conclusion:** Overall, these results demonstrate first proof-of-concept of the potential of CAIX-TAT to treat hypoxic tumors by targeting CAIX-positive hypoxic tumor regions. CAIX-TAT combined with ICI was most effective in inhibiting tumor growth and prolonging survival of tumor-bearing mice. Future studies are required to investigate the radiobiological and immunological effects of CAIX-TAT, to guide optimization of this treatment in combination with ICI.

## Introduction

The tumor microenvironment (TME) is highly important in tumor progression and therapy response. A key feature of the TME is hypoxia, i.e. deprivation of oxygen [1]. Hypoxia can be caused by two main reasons. First, acute hypoxia can occur when blood vessels are temporarily occluded and cells are subjected to a short deprivation of oxygen [2]. Second, chronic hypoxia can occur when cells located further away from blood vessels are subjected to low oxygen levels for prolonged periods of time [3]. In order to survive, chronic hypoxic cancer cells adapt their metabolic and biological state, resulting in a more aggressive and invasive phenotype and contributing to an immunosuppressive TME [4–6]. Furthermore, hypoxic cancer cells are resistant to anti-cancer therapies [4, 6–8], including photon-based external beam radiotherapy (EBRT) and immunotherapy. Therefore, tumor hypoxia is associated with a poor prognosis and remains a major challenge in treating cancer patients.

Carbonic anhydrase IX (CAIX) is an endogenous hypoxia marker upregulated under chronic hypoxia [9, 10]. The potential of CAIX-targeting radiopharmaceuticals for imaging and therapy has been demonstrated in preclinical and clinical studies, as reviewed by Lau et al. [11]. However, these radiopharmaceuticals were mainly tested in renal cell carcinoma models, where CAIX is overexpressed independent of oxygen levels. Recently, [^111^In]In-DTPA-hG250 and [^111^In]In-DTPA-MSC3 have been explored as hypoxia imaging agents in preclinical head-and-neck cancer and melanoma models [12, 13], respectively. These studies showed that anti-CAIX antibodies radiolabeled with imaging radionuclides can target CAIX-expressing hypoxic tumor cells *in vivo*, to identify tumors with a chronic hypoxic phenotype. However, the potential of CAIX-targeting radiopharmaceuticals labeled with therapeutic radionuclides to treat hypoxic tumors has not been investigated yet.

A potentially effective strategy to eliminate hypoxic cancer cells is targeted α therapy (TAT). α-particles, such as actinium-225 (^225^Ac), can cause complex DNA damage independent of oxygen levels, and may therefore overcome the radioresistant phenotype of hypoxic cancer cells [14]. Preclinical studies showed the potential of TAT to eliminate hypoxic tumor cells *in vitro* [15]. However, *in vivo* studies are lacking.

As CAIX-TAT targets hypoxic tumor areas, the normoxic tumor cells are potentially left unaffected. Therefore, combination strategies are required to treat both the normoxic and hypoxic tumor cells. In recent years, immune checkpoint inhibitors (ICI) have proven to be effective anti-cancer agents. By targeting tumor immune escape mechanisms, such as anti-programmed cell death-1 (aPD-1) and anti-cytotoxic T-lymphocyte associated protein 4 (aCTLA-4), T-cell mediated anti-tumor effects can be enhanced. Preclinical studies have shown that targeted radionuclide therapy has immunomodulatory effects and suggest potential for the combination with ICI [16–19]. This is especially of interest for CAIX-TAT, since tumor hypoxia also correlates to immune suppression [6, 13]. Therefore, we hypothesize that CAIX-TAT can eliminate hypoxic tumor cells and that its potential immunomodulatory effects can sensitize tumors to subsequent ICI.

The aim of this study was to investigate the potential of ^225^Ac-labeled anti-CAIX antibody MSC3 ([^225^Ac]Ac-DOTA-MSC3) to treat hypoxic tumors. We determined CAIX-specific binding of radiolabeled MSC3 to B16F10-OVA cells cultured under hypoxic conditions. Subsequently, *in vivo* tumor uptake and spatial correlation to CAIX-expression was determined in B16F10-OVA tumor-bearing mice. Finally, we investigated the therapeutic efficacy of [^225^Ac]Ac-DOTA-MSC3 as monotherapy or in combination with aPD-1/aCTLA-4 ICI in this syngeneic mouse tumor model by measuring tumor growth and survival.

## Materials & methods

### Radiolabeling and cell culture

Details regarding antibody conjugation, radiolabeling and cell culture are described in the supplemental methods.

### Immunoreactive fraction (IRF)

The IRF of [^111^In]In-DOTA-MSC3 and [^225^Ac]Ac-DOTA-MSC3 was determined as previously described [20]. A serial dilution of SK-RC-52 cells (1.1 × 10^6^ – 1.7 × 10^7^ cells/mL, in RPMI (Gibco, Thermo Fisher Scientific, Waltham, MA, USA) containing 0.5% bovine serum albumin (BSA) (Sigma-Aldrich, Saint Louis, MS, USA)) were incubated with 46,729 counts per minute (cpm) [^111^In]In-DOTA-MSC3 (12.4 fmol per well) or 7,326 cpm [^225^Ac]Ac-DOTA-MSC3 (18.8 pmol per well) at 37°C while gently shaken. After 20 min, cells were centrifuged (300 g, 5 min), washed, and the activity in the cell pellet was measured in a γ-counter (Wizard-2480, Perkin Elmer, Waltham, MA, USA). For the quantification of [^225^Ac]Ac-DOTA-MSC3, ^221^Fr was used as a surrogate for ^225^Ac, which is described in more detail in the supplemental methods. The inverse of the specific cell bound activity was plotted against the inverse of the cell concentration in Graphpad Prism (version 5.03), and the IRF was calculated as 1/y-axis intercept.

### Binding and internalization assays

To determine the cell binding and internalization of [^111^In]In-DOTA-MSC3 under hypoxic and normoxic conditions, B16F10-OVA cells were grown in 6-well plates and cultured under hypoxic conditions (1% O_2_) for 24 h in a Whitley H35 Workstation (Don Whitley Scientific) to induce CAIX expression. Cells were washed with phosphate-buffered saline (PBS) and incubated with 36,496 cpm [^111^In]In-DOTA-MSC3 (15.8 fmol per well) in RPMI containing 0.5% BSA at 37°C for 4 h while gently shaken. After incubation, the supernatant was removed, cells were washed twice with PBS, and the cell membrane-bound fraction was collected by incubating the cells for 10 min on ice with acid wash (0.1 M acetic acid, 154 mM NaCl (both Merck Millipore, Burlington, MA, USA), pH 2.6). Subsequently, cells were washed twice with PBS and the internalized fraction was collected by lysing the cells with 1 mL of 0.1 M NaOH (Sigma-Aldrich). Activity in both cell membrane-bound and internalized fractions were counted in a γ-counter.

In a separate experiment, we determined the CAIX-specific membrane binding and internalization. B16F10-OVA cells were grown in 6-well plates and cultured under hypoxic conditions (1% O_2_) for 48 h. Cells were washed with PBS and incubated with 54,069 cpm [^111^In]In-DOTA-MSC3 (22.5 fmol per well) without and with excess of DOTA-MSC3 (110 pmol) in RPMI containing 0.5% BSA at 37°C for 2 h while gently shaken. The membrane-bound and internalized fraction was determined as described previously.

### IC_50_ assay

B16F10-OVA cells were grown in 6-well plates and cultured under hypoxic conditions (1% O_2_) for 24 h as described above. Subsequently, cells were washed with PBS and incubated with increasing concentrations (0.03 – 10.000 nM) of unlabeled DOTA-MSC3 together with 53,370 cpm [^111^In]In-DOTA-MSC3 (13.0 fmol per well) in RPMI containing 0.5% BSA for 4 h on ice. After incubation, cells were washed with PBS and lysed using 1 mL of 0.1 M NaOH. The cell-associated activity was measured using a γ-counter. The IC_50_ value (the concentration of DOTA-MSC3 at which 50% of [^111^In]In-DOTA-MSC3 binding was inhibited) was determined by non-linear regression using one site competition in GraphPad Prism. The graph was normalized by setting the binding of the 0.03 nM unlabeled DOTA-MSC3 condition to 100%.

### Animal experiments

Detailed animal procedures are described in supplemental methods. All animal experiments were conducted in accordance with the principles laid out by the Dutch Act on Animal Experiments. Ethical approval was provided by the institutional Animal Welfare Committee of the Radboud University Medical Center and the Central Authority for Scientific Procedures on Animals (CCD: AVD10300 2020 9645). Experiments were carried out under the supervision of the local Animal Welfare Body. Protocols for each experiment were pre-registered at preclinicaltrials.eu (PCTE0000490, PCTE0000491, PCTE0000589). In total, 79 female wild type C57BL/6J mice (8-10 weeks, Charles River, France) were used. Mice were inoculated subcutaneously with B16F10-OVA cells (1.0×10^6^ cells in 100 µL PBS) on the right flank. Mice were sacrificed by cervical dislocation when humane endpoints (HEPs) were reached or at the end of the experiment as assessed by a blinded biotechnician. HEP were defined as follows: tumor volume > 2.0 cm3, tumor ulceration, >20% body weight loss of their initial body weight, or severe clinical deterioration.

### Biodistribution of [^111^In]In-DOTA-MSC3 at day 7 post injection

The primary outcome of this study was the uptake of radiolabeled DOTA-MSC3 in tumor and the spatial co-localization of radiolabeled DOTA-MSC3 with CAIX-expression. Mice (n=4/group) were included when tumors reached a mean size of 122.9 ± 66.2 mm^3^ (day 11 post tumor cell inoculation). Since it was unknown whether the ^225^Ac signal could be detected with autoradiography, two groups of mice were included. The first group was injected intravenously in the tail vein (i.v.) with 1.2 MBq [^111^In]In-DOTA-MSC3 (13.3 pmol, 90 MBq/nmol plus 346.7 pmol non-labeled DOTA-MSC3) in 200 µl PBS. The second group received 13 kBq [^225^Ac]Ac-DOTA-MSC3 (346.7 pmol, 37.5 kBq/nmol) plus 1.2 MBq [^111^In]In-DOTA-MSC3 (13.3 pmol, 90 MBq/nmol) in 200 µl PBS. Radiochemical purity of [^225^Ac]Ac-DOTA-MSC3 was 81%. Prior to cervical dislocation on day 7 post radiopharmaceutical injections (p.i.), mice were injected with pimonidazole (intraperitoneally (i.p.), 1 h before sacrificing, 2 mg in 500 µL PBS, J.A. Raleigh, Department of Radiation Oncology, University of North Carolina) and with Hoechst 33342 (i.v., 1 min before sacrificing, 0.375 mg in 100 µL PBS, Sigma-Aldrich, Saint Louis, MS, USA). Tumor and organs were dissected. Half of the tumor was formalin-fixed and the other half was snap-frozen in tissue-tek (Sakura, Torrance, CA, USA) using isopentane at −79 °C and used for autoradiography and immunofluorescence (IF) staining. Organs and the formalin-fixed half of the tumors were weighed and the ^111^In signal was measured in a γ-counter (Wizard-2480, Perkin Elmer, Waltham, MA, USA). Radioactivity concentrations were calculated as the percentage of the injected activity per gram of tissue (%IA/g). Two mice (n=1 per group) were excluded from analysis since they reached a HEP (tumor ulceration and an unexpected death) before the end of the experiment.

### Spatial correlation of autoradiography and CAIX IF

To assess the spatial correlation between the intratumoral distribution of radiolabeled MSC3 and CAIX expression, autoradiography and CAIX IF were obtained as described in supplemental methods. Both images were overlayed and analyzed as described previously [21]. In short, IF grey scale images were bicubically rescaled to match autoradiography images in iVision (BioVision Technologies, Exton. PA, USA). Autoradiography images were inverted and overlaid using ImageJ [22]. Parametric mapping of these images was performed in 20 x 20 pixels and gray values were compared for spatial correlation. The gray values within these spatial pixels were correlated based on their intensity in Graphpad Prism, resulting in a R-value.

### Micro-Single Photon Emission Computed Tomography (SPECT)/CT imaging

Uptake of radiolabeled DOTA-MSC3 in CAIX-positive (CAIX+) tumor areas over time was quantified with micro-SPECT/CT imaging. Mice were included when tumors reached a size of 66.3 ± 13.8 mm^3^ (day 7 post tumor cell inoculation). B16F10-OVA tumor-bearing mice (n=8) received an i.v. injection of [^111^In]In-DOTA-MSC3 (19.2 ± 0.2 MBq, 32 µg). Mice were anesthetized with isoflurane, and the torso area was scanned in prone position at 1, 2, 3 and 5 days p.i. using the U-SPECT6/CT MILabs, GP-M0.60 mm multipinhole mouse collimator and MILabs U-SPECT/CT Acquisition software v12.29-st. SPECT scans were acquired in one frame, 20 bed positions using an acquisition time of 12 min (day 1), 15.4 min (day 2), 19.7 min (day 3) and 32.3 min (day 5) per scan. CT scans were acquired for 1.6 min using a one bed position total body scan, default setting with 50 kV and 0.4 mA. At day 5, mice were injected with pimonidazole and Hoechst 33342 as described above and sacrificed by cervical dislocation. Tumors in tissue-tek were snap-frozen and stored at −80 °C. Details regarding SPECT quantification, absorbed dose estimation and the generation of the dose-distribution map are described in supplemental methods.

### Therapeutic efficacy of [^225^Ac]Ac-DOTA-MSC3 monotherapy and in combination with ICI

The primary outcome was tumor growth, expressed as time in days to reach a sixfold tumor volume relative to day 1 (day 1 was selected since at day 0 the tumor volume measurements are less accurate). For each mouse, the tumor size at day 1 was set at 100% and the tumor measurements were fit by an exponential fit or a one-phase association fit (constrain: Y0=100). In one case, tumor reduction was followed by growth; therefore, only the growth phase was fitted exponentially. Fits with an R^2^-value above 0.5 were considered in the analysis. Fitted curves reaching a plateau below a sixfold tumor volume, were excluded for the analysis. The secondary outcome was Kaplan-Meier survival with the HEP considered as survival endpoint. At the start of the experiment, 10 mice/group were included when B16F10-OVA tumors reached a size of 67.0 ± 22.4 mm^3^ (day 8). The following treatment regimens were studied: 1) PBS, 2) unlabeled DOTA-MSC3, 3) ICI, 4) [^225^Ac]Ac-DOTA-MSC3, 5) [^225^Ac]Ac-DOTA-MSC3 + ICI (Figure 1). [^225^Ac]Ac-DOTA-MSC3 treatment consisted of a i.v. injection of 15 kBq [^225^Ac]Ac-DOTA-MSC3 (30 µg) in 200 µl PBS at day 0. ICI treatment consisted of an i.p. injection of aPD-1/aCTLA-4 (200 µg of BE0146-clone RMP1-14 and of BE0032-clone UC10-4F10-11 (both Bio X Cell, Lebanon, NH, USA)) in 200 µl PBS at day 1, 4, 7, 10, 13 and 16. Control groups received an i.v. injection of PBS (200 µl) or 30 µg non-labeled MSC3 in 200 µl PBS (both at day 0) and i.p. injection of PBS (200 µl at day 1, 4, 7, 10, 13 and 16). Tumor volume, survival and body weight were monitored until the end of follow-up, predefined at 6 weeks post start of treatment.

**Figure 1:**
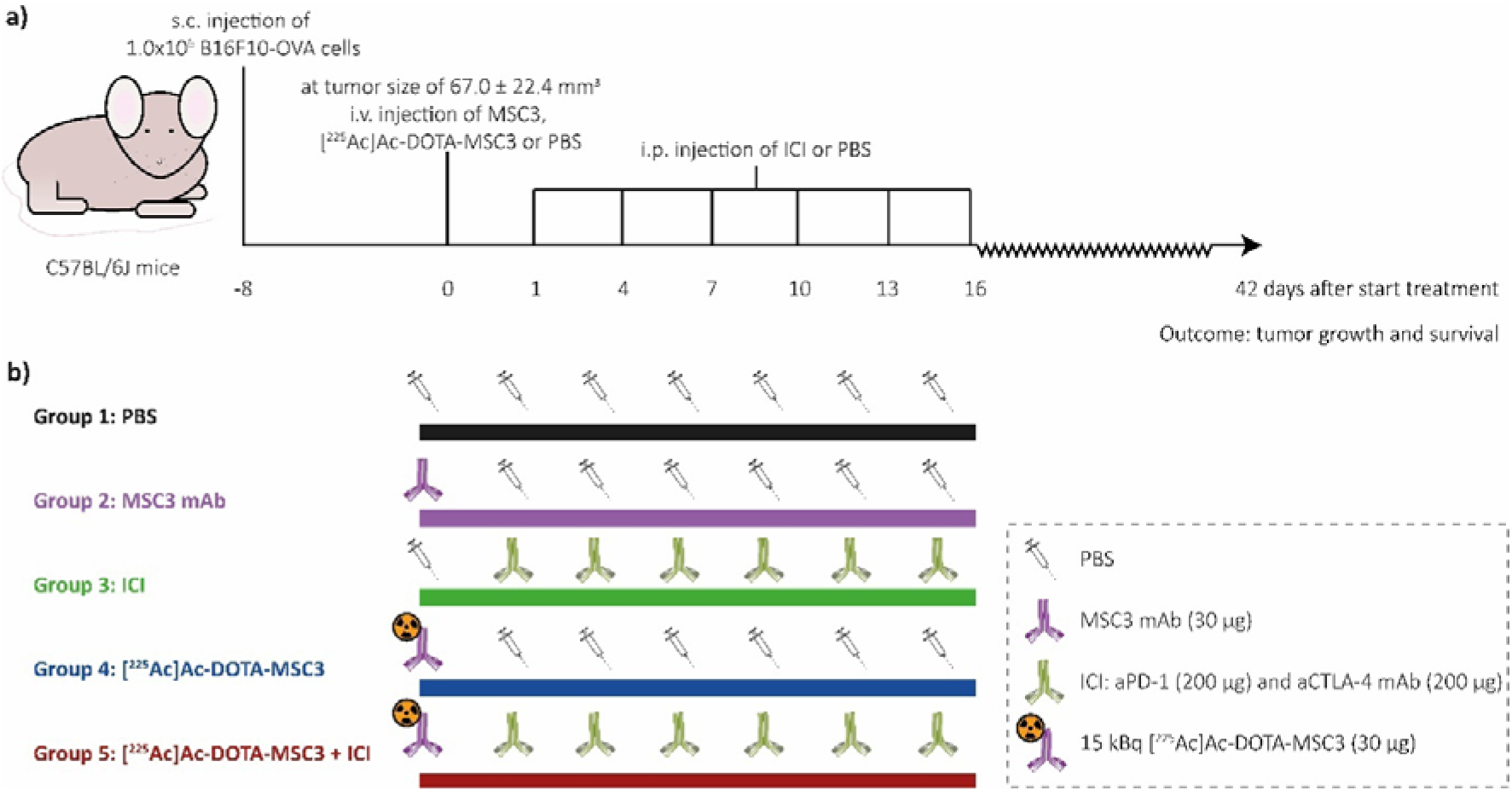
Schematic overview of the therapeutic efficacy study **a)** General experimental set-up. Mice were followed until 42 days p.i. or until HEP were reached **b)** Overview of the experimental groups (n=10 mice/group).

### Ex vivo CD45 IF and quantification

To investigate immune cell infiltration upon [^225^Ac]Ac-DOTA-MSC3, ICI and combination therapy, CD45 staining and quantification was performed on B16F10-OV tumor sections at 5 and 8 days post treatment. At the start of the experiment, 7 mice per group were included (mean B16F10-OVA tumor size of 49.0 ± 21.7 mm3 (day 8 post tumor cell inoculation)). The a priori sample size calculation was based on the expected treatment effect as estimated using quantification of pilot tumor material, using a significance level of 5%, a power of 80% and a two-way ANOVA. The following treatment regimens were studied: 1) Baseline control at day 0; 2) PBS control day 5 and 8; 3) ICI day 5 and 8; 4) [^225^Ac]Ac-DOTA-MSC3 day 5 and 8; 5) [^225^Ac]Ac-DOTA-MSC3 + ICI day 5 and 8.

Treatment dose and schedules were same as ascribed for the therapeutic efficacy study above. In total, seven mice (listed in Table S1) were excluded from analysis since they reached a HEP (tumor ulceration and or severe clinical deterioration) before the end of the experiment. Tumors were collected at 5 and 8 days post treatment. B16F10-OVA FFPE tumor sections (4 µm) were mounted on FLEX IHC Microscope Slides, deparaffinated and dehydrated, followed by antigen retrieval in 10 mM sodium citrate pH 6.0. Between consecutive steps of the staining process, sections were rinsed 3 times in Tris-buffered saline with Tween (TBST, Merck, NJ, USA). Endogenous peroxidase activity was blocked at RT by 10 min incubation with 3% H_2_O_2_ (Merck) and non-specific binding was blocked by 10 min incubation with TBST + 1% BSA (Sigma-Aldrich, Saint Louis, MS, USA). Immediately after, sections were incubated at RT for 1h with primary anti-CD45 antibody (1:150 dilution in in TBST + 1% BSA, #70257, Cell Signaling Technology). Slides were incubated with secondary antibody at RT by 30 min incubation with Opal polymer anti-mouse with anti-rabbit HRP (#ARH1001EA, Akoya Biosciences, MA, USA), followed by Opal-620 fluorophore working solution (1:200, #FP1495001KT, Akoya Biosciences, MA, USA) according to manufacturer’s instructions. Antigen retrieval was performed with 10 mM sodium citrate. Slides were incubated at RT for 2 min with DAPI (#D1306, ThermoFisher, MA, USA) and mounted a cover slip using Fluoromount (F2680 Sigma-Aldrich). For each tumor section, grey scale images (pixel size of 0.64 X 0.64 µm) of nuclei and CD45 staining were obtained (20 random selected images each, for smaller tumors >10 each). Tumors sections with <10 snapshots were excluded (reasons listed in Table S1). Grey scale images were converted into binary images. For nuclei staining, binary images were generated manually above the background staining. Binary images were used to quantify the mean CD45 area relative to the tumor area. For each group, the means of each mice was used for analysis.

### Statistics

Differences in uptake in tumor or organs between mice injected with [^111^In]In-DOTA-MSC3 alone or in combination with [^225^Ac]Ac-DOTA-MSC3 were compared using multiple unpaired t-tests with Holm-Šídák multiple comparisons tests (Table S2). In the SPECT quantification study, differences in CAIX+ tumor volumes and differences in the CAIX+ and CAIX-tumor uptake between different time points were compared using an one-way ANOVA Kruskal-Wallis test. In the therapy study, analyses entailed comparison of treatment groups versus PBS and the combination therapy group versus ICI and [^225^Ac]Ac-DOTA-MSC3 monotherapy. Tumor growth was analyzed by comparing differences in time to reach a sixfold tumor volume relative to day 1 (Table S3) and by comparing normalized tumor volume over life-time of mice area under the curve (Figure S1) using Kruskal-Wallis Test with Dunn’s multiple comparisons test. Mean tumor volumes at day 1 were compared using Kruskal-Wallis Test with Dunn’s multiple comparisons test (Figure S2). Kaplan-Meier survival plots of the groups were obtained and were compared using pairwise Mantel-Cox log-rank tests and Bonferroni correction (Table S4). Differences in CD45-positive tumor areas was executed between groups were tested using an one-way ANOVA Šidák’s multiple comparisons test. Significance levels (α) of 0.05 were used or the Bonferroni-corrected α for survival comparisons.

## Results

### 111In- and ^225^Ac-labeled DOTA-MSC3 specifically bind to CAIX-expressing hypoxic cancer cells

[^225^Ac]Ac-DOTA-MSC3 was obtained with high stability for up to 11 days (Figure 2a). In vitro binding assays demonstrated a high IRF for [^111^In]In-DOTA-MSC3 (94%, Figure 2b) and [^225^Ac]Ac-DOTA-MSC3 (90%, Figure 2c). B16F10-OVA cells were cultured under hypoxic conditions to induce CAIX expression [13]. [^111^In]In-DOTA-MSC3 showed increased binding to and internalization in B16F10-OVA cells when cultured under hypoxic versus normoxic conditions (Figure 2d). This binding was partially blocked by adding an excess unlabeled DOTA-MSC3, indicating CAIX specificity under hypoxic culturing conditions (Figure 2e). DOTA-MSC3 bound with high affinity to hypoxic B16F10-OVA cells (IC_50_ = 3.6 nM, Figure 2f).

**Figure 2:**
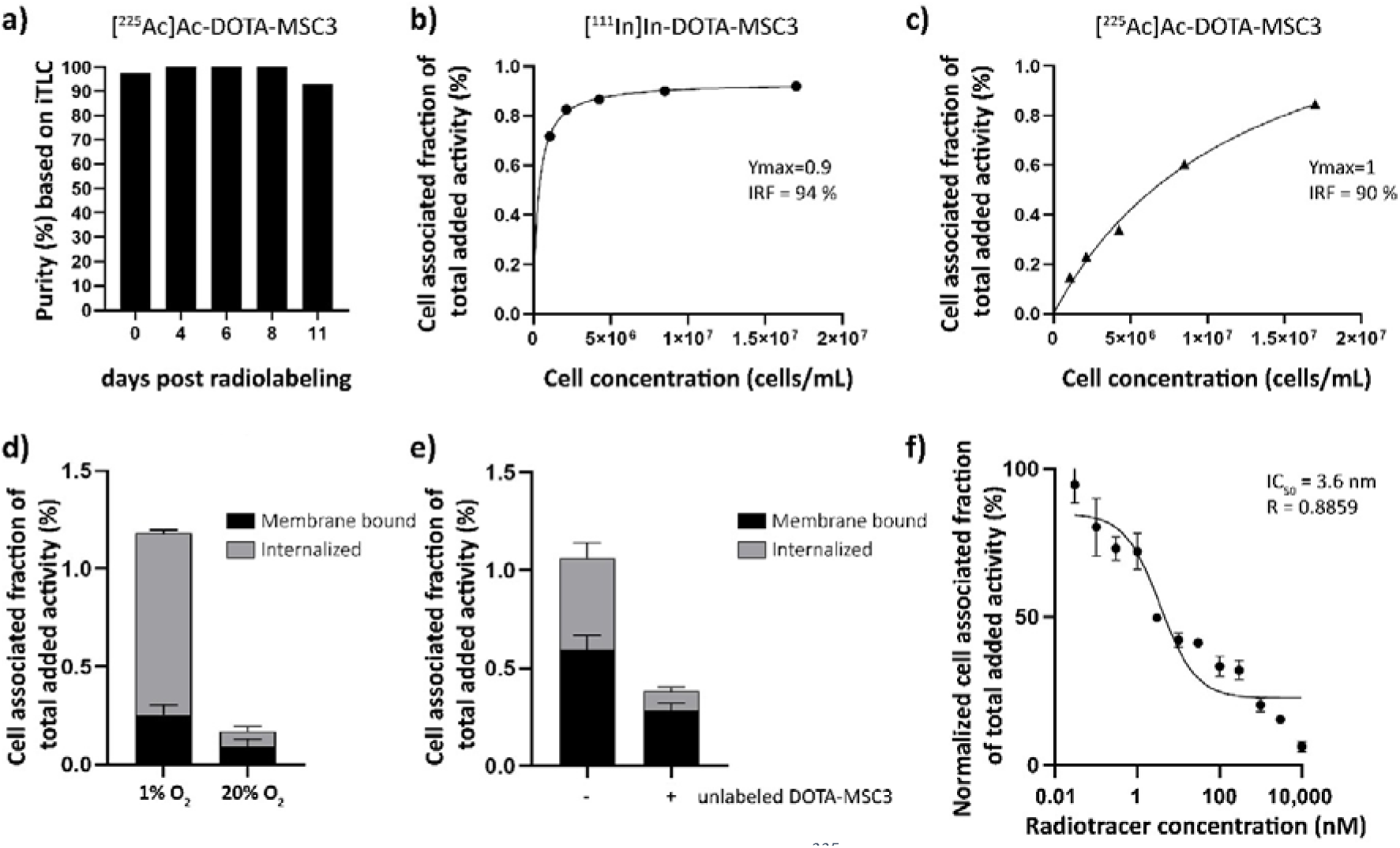
In vitro characteristics of (radiolabeled) MSC3 **a)** Stability of [^225^Ac]Ac-DOTA-MSC3 after 4, 6, 8 and 11 days post radiolabeling determined by iTLC (n=1), **b and c)** Lindmo assays in SK-RC-52 cells. **d)** Binding of [^111^In]In-DOTA-MSC3 in B16F10-OVA cells cultured in hypoxic conditions or normoxic conditions and **e)** without and with presence of a molar excess of unlabeled DOTA-MSC3, **f)** IC_50_ assay of DOTA-MSC3. SD represents technical replicates.

### 111In- and ^225^Ac-labeled DOTA-MSC3 accumulate in CAIX-expressing hypoxic tumor regions in vivo

To determine the uptake and intratumoral distribution of radiolabeled DOTA-MSC3, mice were injected with [^111^In]In-DOTA-MSC3 alone or co-injected with [^225^Ac]Ac-DOTA-MSC3. The ex vivo biodistribution at 7 days p.i. was quantified by γ-counting of ^111^In (Table S2). Tumor uptake was 13.0 ± 4.4 %IA/g in mice that received [^111^In]In-DOTA-MSC3, and 16.1 ± 3.5 %IA/g in mice that were co-injected with [^225^Ac]Ac-DOTA-MSC3 (p=0.9694). This shows that co-injection with [^225^Ac]Ac-DOTA-MSC3 did not affect the tumor uptake of [^111^In]In-DOTA-MSC3. Autoradiography visualized heterogeneous distribution of ^111^In- and ^225^Ac-labeled DOTA-MSC3 in B16F10-OVA tumors (Figure 3a and b). The ^225^Ac-derived autoradiography signal was much higher compared with the ^111^In-derived signal. This indicates that the phosphor imager plate was activated more efficiently by the high-energy emissions of ^225^Ac and its daughters, compared with the relatively low-energy γ-emission of ^111^In. The radiosignal co-localized with CAIX-expression and pimonidazole (hypoxia) staining on consecutive slides with a high spatial correlation of R=0.47 ± 0.13 (Figure S3). Extended IF images (CAIX, pimonidazole, Hoechst, 9F1) are shown in Figure S4.

**Figure 3:**
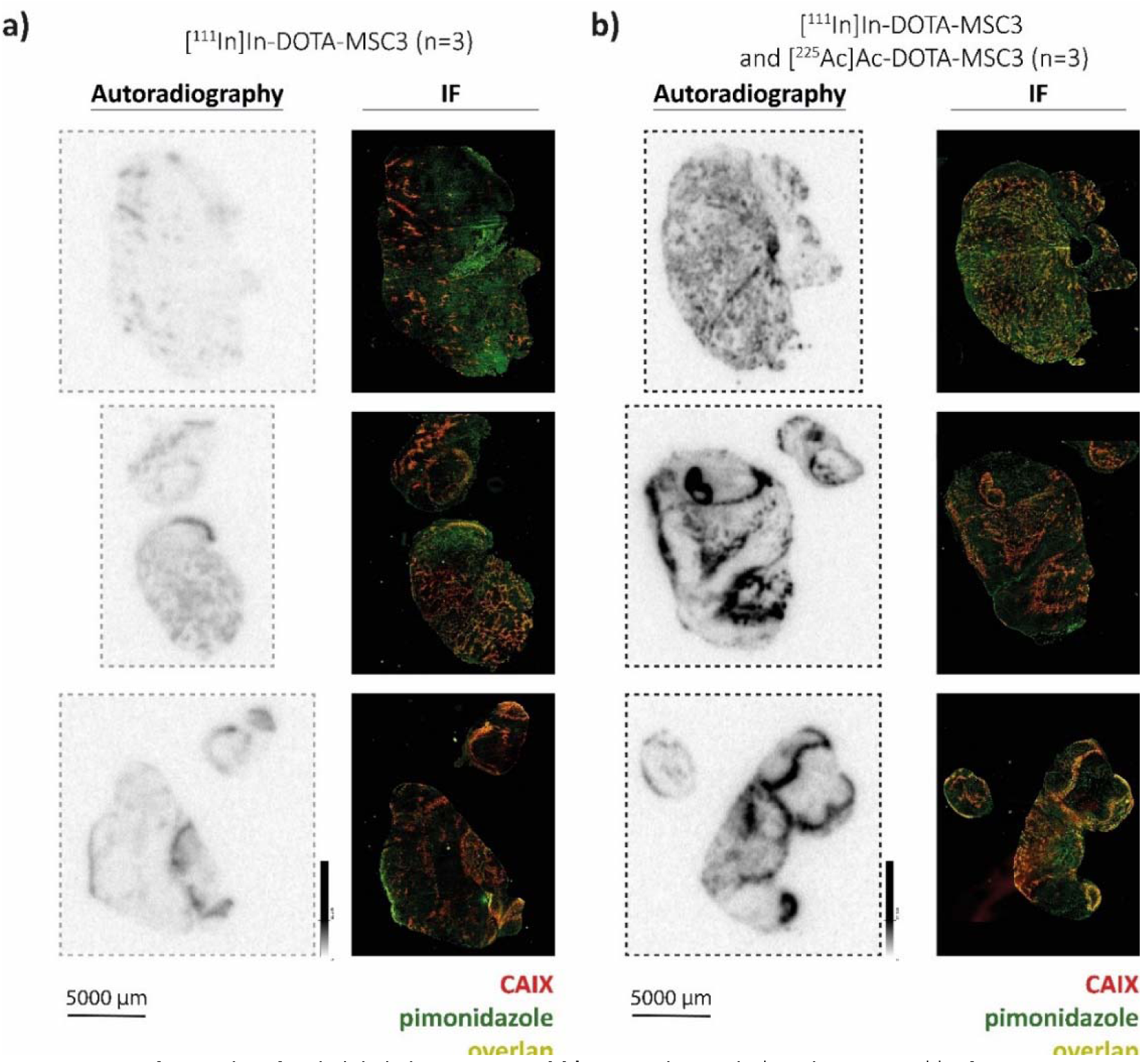
CAIX-specific uptake of radiolabeled MSC3. **a and b)** Autoradiography* and IF images** of B16F10-OVA tumor sections (n=3/group). *Autoradiography images display unaltered signal intensities to allow direct visual comparison, **Brightness of IF images are adjusted for visualization purposes, the original images are used for co-registration analysis.

### CAIX-specific uptake and retention of [^111^In]In-DOTA-MSC3 in vivo

The uptake of [^111^In]In-DOTA-MSC3 in CAIX+ tumor areas over time was determined based on Micro-SPECT/CT imaging. Figure 4a shows a representative scan at 5 days p.i. (additional scans are presented in Figure S5). The mean CAIX+ tumor volume of the total tumor volume was 13.7 ± 8.4 % on day 1 p.i. and shows a trend towards reduction over time (7.5 ± 4.1 % on day 2 p.i.), although this was not statistically significant (Figure 4b). Uptake of [^111^In]In-DOTA-MSC3 in the CAIX+ regions was approximately five times higher than in CAIX-negative (CAIX-) regions (23.1 ± 4.8 %IA/mL versus 4.4 ± 1.1 %IA/mL on day 5, Figure 4c). The spatial correlation between the radiosignal and CAIX expression was confirmed for tumors harvested at 5 days p.i. (R=0.53 ± 0.12, example shown in Figure 4e).

**Figure 4:**
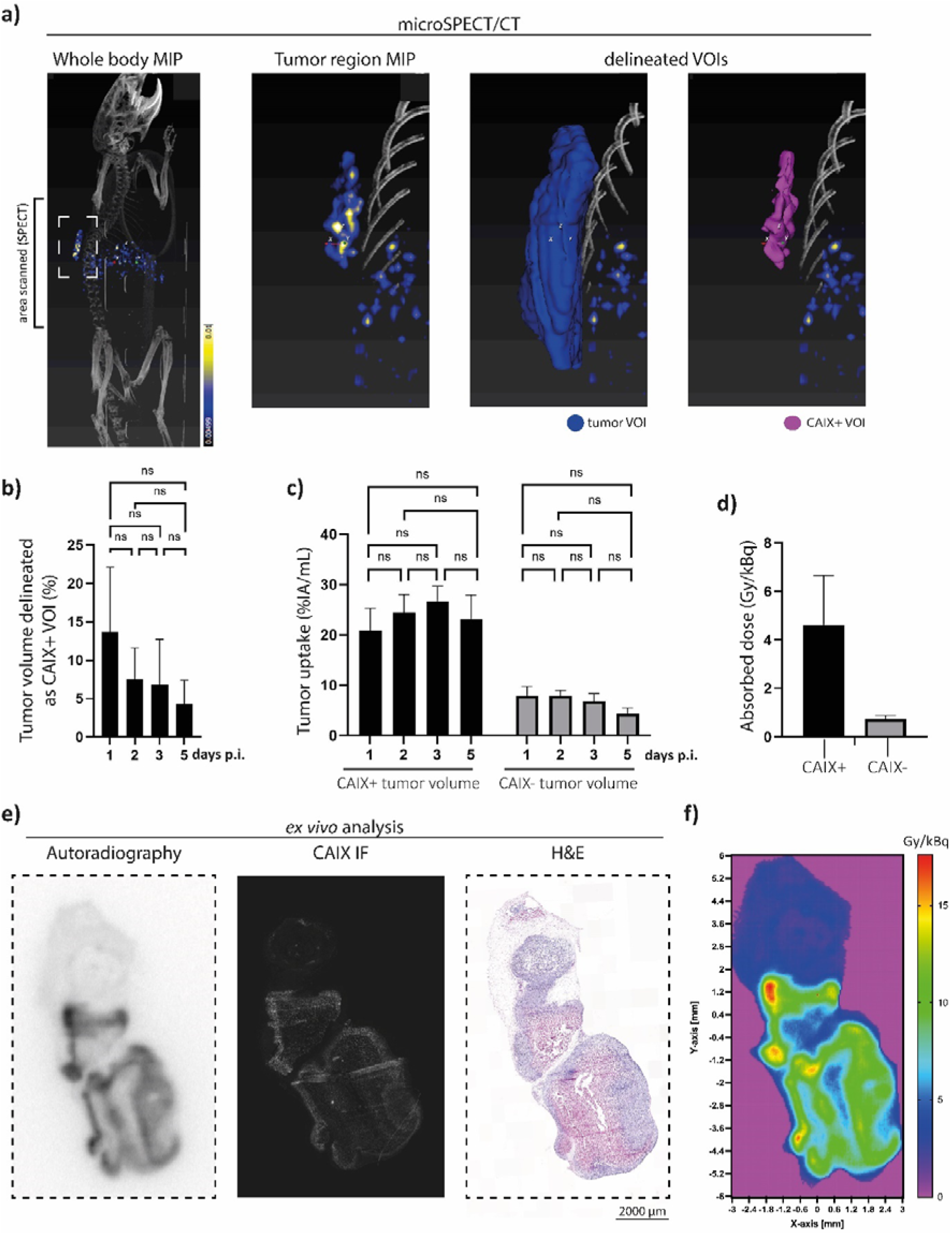
Accumulation of [^111^In]In-DOTA-MSC3 in B16F10-OVA tumors a) A representative SPECT/CT image (torso are**a)** at 5 days p.i. and delineated volume of interest (VOIs), **b)** CAIX+ tumor volumes in percentage of the total tumor volume over time, **c)** Uptake over time, **d)** Absorbed dose in CAIX+ and CAIX-tumor volumes, **e)** example of ex vivo autoradiography, CAIX IF and Hematoxylin and Eosin (H&E) staining **f)** Absorbed dose map of ^225^Ac. SD represents variation between all mice (n=7).

### Estimated absorbed tumor dose for [^225^Ac]Ac-DOTA-MSC3

Dosimetry calculations predicted a six times higher absorbed dose for the CAIX+ tumor volume (4.63 ± 2.0 Gy/kBq) compared with the CAIX-tumor volume (0.73 ± 0.15 Gy/kBq) for [^225^Ac]Ac-DOTA-MSC3 (Figure 4d). Additionally, the dose-distribution autoradiography map of ^225^Ac shows a heterogenous dose-distribution within the tumor (Figure 4f).

### Treatment with [^225^Ac]Ac-DOTA-MSC3 plus ICI prolongs survival and delays tumor growth in B16F10-OVA tumor-bearing mice

To determine therapeutic efficacy, B16F10-OVA tumor-bearing mice were injected with [^225^Ac]Ac-DOTA-MSC3 with or without ICI. The activity dose of 15 kBq was selected to reach an estimated absorbed dose of 69.5 ± 30.0 Gy for the CAIX+ tumor volume. [^225^Ac]Ac-DOTA-MSC3 seemed to delay tumor growth, although the time to reach a sixfold tumor volume relative to day 1 did not differ significantly from PBS control (Figure 5a and b). However, the combination of [^225^Ac]Ac-DOTA-MSC3 plus ICI did significantly inhibit tumor growth as compared with PBS (p=0.0022) and ICI (p=0.0048). ICI and MSC3 mAb did not alter tumor growth compared with PBS. The median survival of mice that received [^225^Ac]Ac-DOTA-MSC3 monotherapy or in combination with ICI were 18 days and 21 days, respectively. The median survival of mice that received PBS, non-labeled MSC3 or ICI as control, were 9.5, 8 and 8.5 days, respectively. Survival upon [^225^Ac]Ac-DOTA-MSC3, both alone and in combination with ICI, was significantly longer as compared with PBS (Figure 5c). Body weight was relatively stable over time (Figure S6). Some mice lost weight, but this did not exceed the predefined HEP criteria. Overall, several mice (78%) reached HEP due to severe clinical deterioration, which was observed across all treatment and control groups (Figure S7).

**Figure 5:**
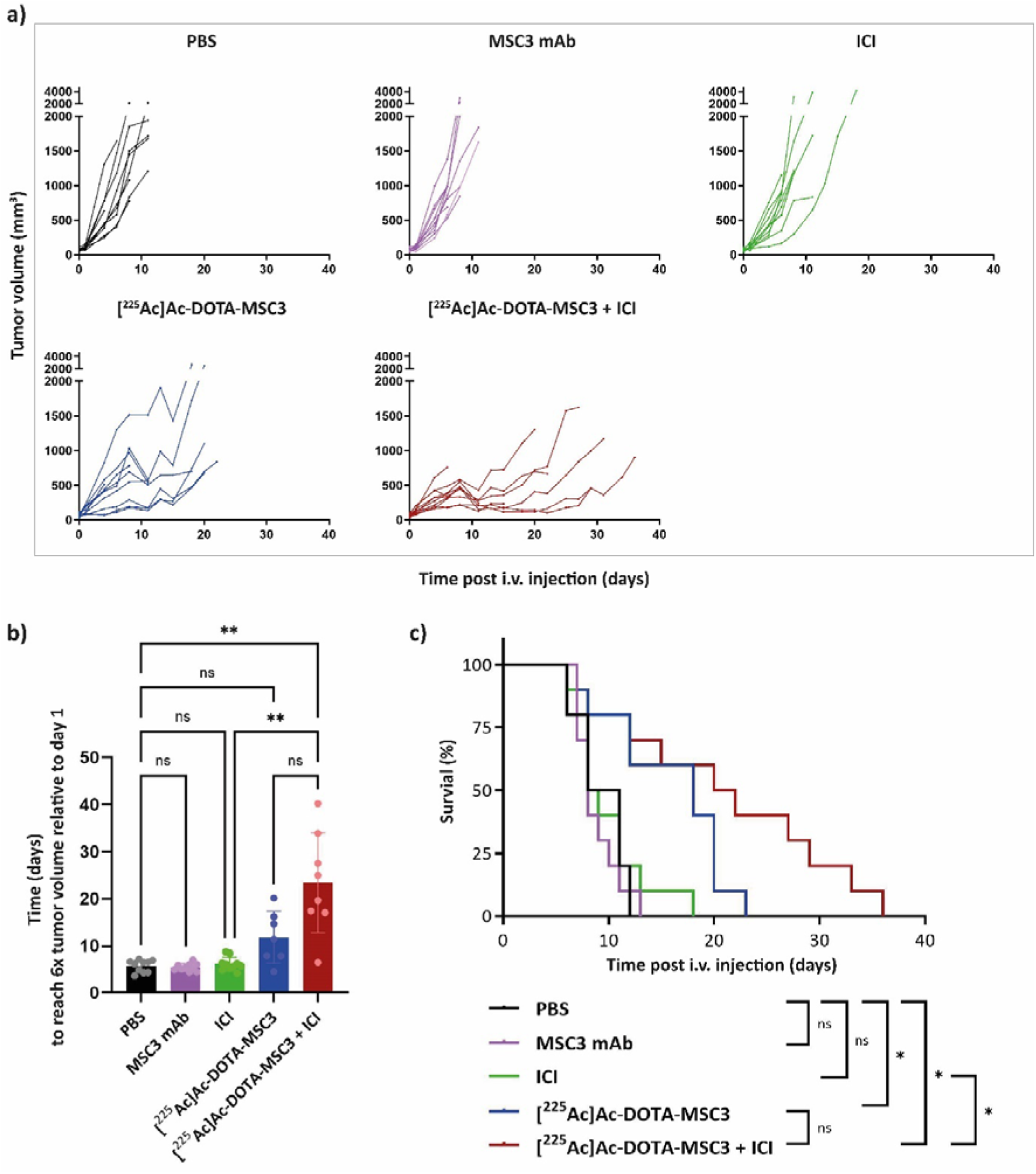
Therapeutic effect of [^225^Ac]Ac-DOTA-MSC3 with or without ICI in B16F10-OVA tumor-bearing mice (n=10/group) **a)** Tumor growth curves of individual mice **b)** Time in days to reach 6x tumor volume relative to day 1 c) Kaplan-Meier survival. SD represents variation between all mice.

### Immune infiltration after [^225^Ac]Ac-DOTA-MSC3 and ICI treatment

Presence of CD45-expressing cells was determined by calculating the fraction of tumor area that was positive for CD45 IF. The combination treatment with [^225^Ac]Ac-DOTA-MSC3 and ICI resulted in a significant increase in CD45 positive tumor area, compared with ICI monotherapy 8 days p.i. (p = 0.01) (Figure 6). Although not significant, a trend for higher CD45-positive tumor areas was observed for mice treated with the combination ([^225^Ac]Ac-DOTA-MSC3 and ICI) compared with [^225^Ac]Ac-DOTA-MSC3 monotherapy at day 8 and compared with ICI monotherapy at day 5. No significant differences were shown for [^225^Ac]Ac-DOTA-MSC3 and ICI monotherapies compared to PBS at both time points.

**Figure 6:**
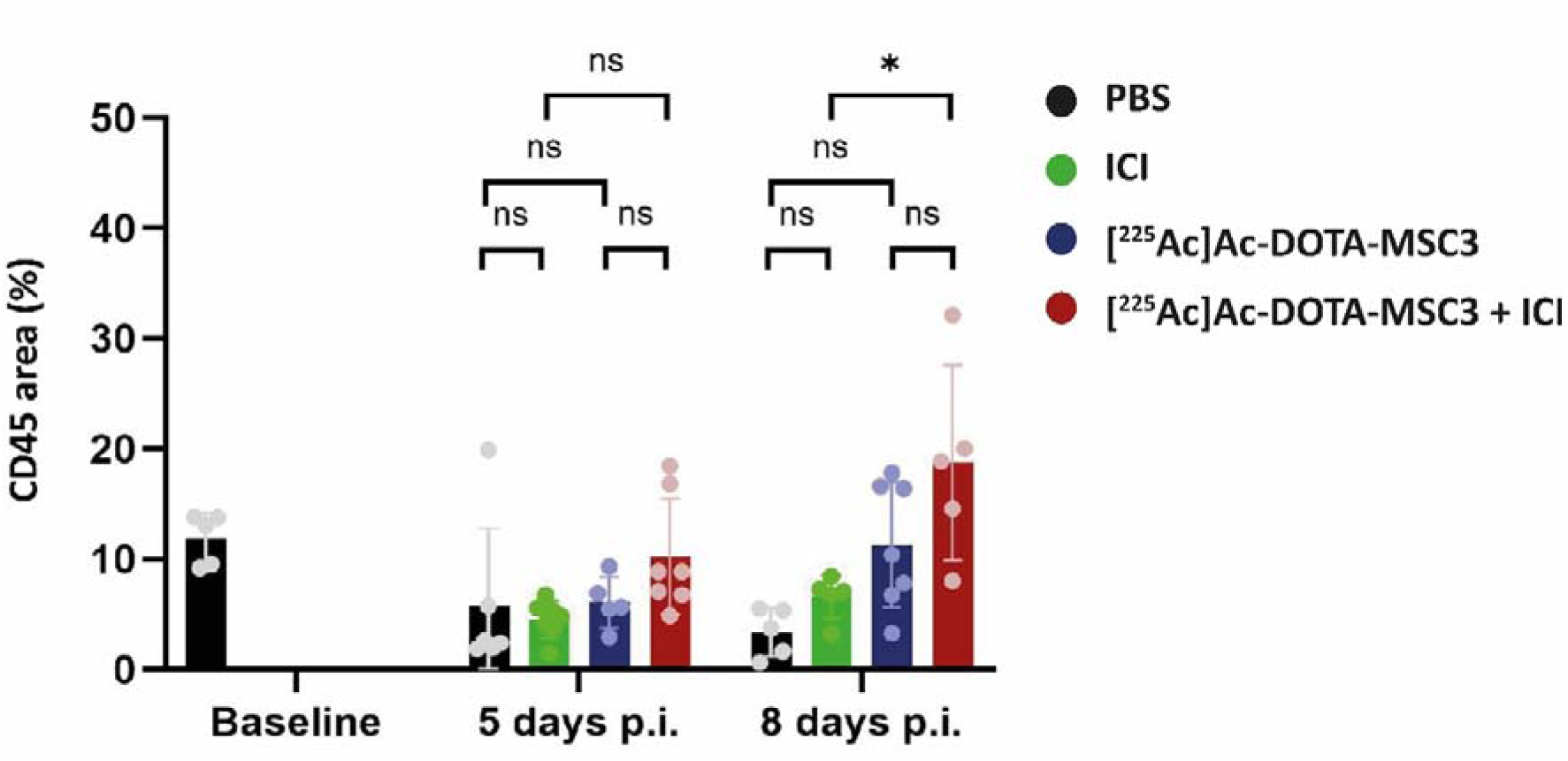
Quantification of CD45 area (%) in B16F10-OVA tumor sections at 5 and 8 days p.i. of untreated mice (baseline), mice treated with PBS (black), ICI (green), [^225^Ac]Ac-DOTA-MSC3 (blue) or [^225^Ac]Ac-DOTA-MSC3 with ICI (red).

## Discussion

Tumor hypoxia remains a major clinical challenge, underlining the need for hypoxia-specific therapeutic interventions. Multiple strategies to overcome tumor hypoxia have been investigated, but were not translated or implemented in the clinic (reviewed by Overgaard et al. [23]). In this study, we demonstrated the tumor targeting and therapeutic potential of ^225^Ac-labeled MSC3 in a hypoxic syngeneic mouse tumor model. We showed that [^225^Ac]Ac-DOTA-MSC3 targets the CAIX+ hypoxic areas and prolongs survival of tumor-bearing mice. As [^225^Ac]Ac-DOTA-MSC3 only targets CAIX+ areas, combination therapies are needed to eliminate the whole tumor. Here, we showed that the combination of [^225^Ac]Ac-DOTA-MSC3 with ICI significantly inhibits tumor growth and prolongs survival.

In vitro, [^225^Ac]Ac-DOTA-MSC3 binds to CAIX with a IRF comparable to other ^225^Ac-labeled CAIX-targeting antibodies [24, 25]. The relatively low binding and uptake of [^225^Ac]Ac-DOTA-MSC3 in B16F10-OVA cells and tumors, compared to SK-RC-52 cells and tumors [13, 25], can be attributed to the fact that SK-RC-52 cells constitutively overexpress CAIX independent of oxygen levels, whereas only ∼8% of the B16F10-OVA tumor expresses CAIX (Figure 4b). The remaining tumor cells express no or only minimal levels of CAIX. Therefore, CAIX-specific tumor uptake is underestimated when considering the total tumor uptake. The intratumoral distribution of ^111^In- and ^225^Ac-labeled DOTA-MSC3 spatially correlated with the expression of CAIX, which is in line with previous data [13]. Interestingly, the autoradiographical analysis of tumor sections from mice that were co-injected with ^111^In- and ^225^Ac-labeled DOTA-MSC3 showed a much higher signal intensity compared with tumor sections from [^111^In]In-DOTA-MSC3 injected mice, despite the lower injected activity (1.2 MBq ^111^In versus 13 kBq ^225^Ac). This suggests that the phosphor imager plate is activated more efficiently by the high energy β-emission during ^225^Ac decay than by the relatively low-energy γ-emission of ^111^In [26]. Notably, in this experiment the radiochemical purity of [^225^Ac]Ac-DOTA-MSC3 was only 81%. This may have resulted in an underestimation of the CAIX-specific uptake in the tumor, while increased uptake may have occurred in organs that accumulate ^225^Ac such as liver and bone [27].

To perform dosimetric calculations and obtain absorbed dose values for [^225^Ac]Ac-DOTA-MSC3, we used [^111^In]In-DOTA-MSC3 as a surrogate radiotracer as this allows sensitive and accurate longitudinal SPECT imaging. The dosimetry results should be interpreted with caution for various reasons. First of all, we assumed comparable biodistribution of ^111^In- and ^225^Ac-labeled DOTA-MSC, however, if this assumption is incorrect, this could lead to incorrect absorbed dose estimations. Secondly, we did not consider the recoil effect of ^225^Ac. Daughter radionuclides can redistribute to other tissues, for example the kidneys [25, 28]. This redistribution may lead to an overestimation of the tumor absorbed dose and an underestimation of the kidney absorbed dose. Finally, the uptake and the absorbed dose of the CAIX+ volume of interest (VOI) may be underestimated due to the partial volume effect (PVE) that can cause blurring of signals from and to the surrounding area, especially for small volumes [29]. A possible solution to correct for this is to use phantoms that represent the geometry of the VOI [30], however small and variable CAIX+ VOIs prevented accurate phantom PVE measurements in this study.

We demonstrated that [^225^Ac]Ac-DOTA-MSC3 significantly prolongs the survival of B16F10-OVA tumor-bearing mice compared with PBS. Additionally, the combination of [^225^Ac]Ac-DOTA-MSC3 with ICI significantly delayed tumor growth and significantly prolonged survival compared with PBS and ICI. A trend might be observed indicating improved survival and delayed tumor growth when comparing the combination of [^225^Ac]Ac-DOTA-MSC3 plus ICI with [^225^Ac]Ac-DOTA-MSC3 monotherapy, however this was not significantly different. In the tumor growth curve analysis, we excluded curves that plateaued at less than the sixfold tumor volume relative to day 1 (n=3 in [^225^Ac]Ac-DOTA-MSC3 group and n=2 in the combination group). As these are the best-responding mice, this analysis underestimates the treatment effect.

The therapeutic potential of CAIX-TAT has been studied previously in other models. Morgan et al. and Merkx et al. demonstrated that ^225^Ac-labeled anti-CAIX antibody hG250 inhibits tumor growth in immunodeficient mice bearing SK-RC-52 tumors [24, 25]. Of note, the unique aspects of our mouse model are 1) tumors that only expresses CAIX under hypoxic conditions, 2) which are grown in fully immunocompetent mice. As we hypothesize that combination treatment strategies will be necessary to eliminate both hypoxic and normoxic tumor cells. In this study we focused on the combination with ICI. However, other combination therapies such as EBRT or chemotherapy could be applied as well.

A limitation of our study is the rapid growth of B16F10-OVA tumors, which is not representative of the human situation and limits long-term follow-up. Biodistribution studies with [^111^In]In-DOTA-MSC3 demonstrated uptake in liver, lung and spleen (in line with previous findings [13]), which may be due to physiological clearance or low CAIX expression [31, 32]. Increased uptake (in %IA/g) in the spleen after co-injection with [^225^Ac]Ac-DOTA-MSC3 may be due to atrophy, as a decline in spleen mass was observed [25]. During the therapy study we evaluated toxicity based on body weight and the general well-being of the animal. Several mice were taken out of the study because of severe clinical deterioration. However, as this occurred across all treatment and control groups this was most likely caused by the tumor rather than by the therapies.

In this study we used a monoclonal antibody for TAT. But as antibodies slowly clear from the circulation, and as the recoil effect could cause redistribution of daughter radionuclides through the body, high absorbed doses in healthy organs can be obtained, potentially leading to toxicity in these organs. To improve the pharmacokinetics, smaller-sized radiopharmaceuticals could be used. In this regard, antibody fragments and nanobodies have been investigated for CAIX-targeting of hypoxic tumor areas [12, 33]. These studies showed rapid clearance, but, at the cost of tumor uptake as compared to the full antibody. Other alternatives are CAIX-targeting small molecules [34] or CAIX-targeting peptides [35]. However, the potential of these novel CAIX radiopharmaceuticals to specifically target hypoxic tumor areas has not been demonstrated yet. Taken together, there is a clear need to develop CAIX-TAT-radiopharmaceuticals with a favorable pharmacokinetic profile combined with high uptake and good retention in hypoxic tumor cells.

Although this study shows first proof-of-concept of CAIX-TAT to treat hypoxic tumor cells, some open questions remain, especially on the radiobiological and immunological effects. We have shown first indications that [^225^Ac]Ac-DOTA-MSC3 combined with ICI results in an increased infiltration of CD45-positive immune cells, compared with ICI alone. However, more in depth characterization of the subsets of immune cells that are primarily involved is warranted. Furthermore, a direct comparison of CAIX-TAT with EBRT or TRT with beta-emitting radionuclides has not been performed yet. Finally, different [^225^Ac]Ac-DOTA-MSC3 doses, ICI doses and administration schedules could result in more effective treatment regimens. Future research addressing these questions would provide more knowledge on the mechanism-of-action of CAIX-TAT and can inform on optimal timing and treatment schedules for clinical translation, or identify which tumors or patient populations may benefit from this therapy.

### Conclusion

This study demonstrates first proof-of-concept of the potential of CAIX-TAT to treat hypoxic tumors. The combination of [^225^Ac]Ac-DOTA-MSC3 with ICI was most effective in inhibiting tumor growth and prolonging survival of B16F10-OVA tumor-bearing mice. Future studies are required to investigate the radiobiological and immunological effects of CAIX-TAT in hypoxic and normoxic tumor regions, to guide optimization of this treatment in combination with ICI.

## Abbreviations

TAT: Targeted α therapy; ICI: immune checkpoint inhibitors; ^111^In: indium-111; ^225^Ac: actinium-225; TME: tumor microenvironment; EBRT: external beam radiotherapy; CAIX: carbonic anhydrase IX; aPD-1: anti-programmed cell death-1; aCTLA-4: anti-cytotoxic T-lymphocyte associated protein 4; IRF: immunoreactive fraction; cpm: counts per minute; PBS: phosphate-buffered saline; HEP: humane endpoint; IF: immunofluorescence; i.v.: injected intravenously in the tail vein; p.i.: post radiopharmaceutical injections; i.p.: intraperitoneally; %IA/g: percentage of the injected activity per gram of tissue; CAIX+: CAIX-positive; CAIX-: CAIX-negative; SPECT: Micro-Single Photon Emission Computed Tomography; SD: standard deviation; VOI: volume of interest; H&E: Hematoxylin and Eosin.

## Supplementary Material

Supplementary methods, figures and tables are provided in the supplementals.

## Acknowledgements

We thank Rowan Wuestenenk, Milou Boswinkel, Cathelijne Frielink, Paul Rijken and Luuk Oostveen for their valuable discussions and practical assistance. We thank Floor Moonen, Liz van den Brand, Mariëlle van de Berkt-van Ginkel, Alex Hanssen, Bianca Lemmers-van de Weem, Karin de Haas-Cremers and Kitty Lemmens-Hermans for technical assistance with the animal experiments. All animal experiments were performed at the Radboud Preclinical Imaging Center (PRIME). We thank MiLabs for assistance in the SPECT quantification. Parts of the study results have been presented at the 12^th^ International Symposium on Targeted Alpha Therapy 2023 conference, the 2^nd^ workshop on Radiobiology of Molecular Radiotherapy 2023 conference, the European Society for Molecular Imaging conference in 2024, the European Association of Nuclear Medicine conference in 2024 and the International Workshop on Radiobiology of Molecular Radionuclide Therapy in 2025 and published in the Abstract books.

## Author Contributions

All authors contributed to the study conception and design. Material preparation, data collection and analysis were performed by Sylvia Wenker, Mark Konijnenberg, Daphne Lobeek, Giulia Tamborino, Janneke Molkenboer-Kuenen, Gerben Franssen, Daan Boreel, Simone Kleinendorst and Hans Peters. The first draft of the manuscript was written by Sylvia Wenker, Johan Bussink, Sanne van Lith and Sandra Heskamp and all authors commented on previous versions of the manuscript. All authors read and approved the final manuscript.

## Data availability

Data will be made available on request to the corresponding author.

