## Supplementals for "CAIX-targeted α therapy directed against hypoxic tumor cells in combination with immune checkpoint inhibitors in a syngeneic mouse tumor model"

### **Supplemental Materials & Methods**

#### *Conjugation*

Anti-murine and human CAIX mAb (clone: MSC3) was obtained from Creative Biolabs (TAB-1458CL, New York, USA) and functionalized with p-SCN-Bn-DOTA (1020407-41-3 Macrocyclics, Plano, USA) as described previously [1]. In short, MSC3 was concentrated and dialyzed against phosphate-buffered saline (PBS) using Amicon Ultra-4 20,000 MWCO (Merck Millipore, Burlington, MA, USA). P-SCN-Bn-DOTA (15 eq) in DMSO (Merck) was added to concentrated MSC3 in 0.1 M NaHCO_3_ buffer (pH=9) (Sigma-Aldrich, Saint Louis, MS, USA) and shaken at 350 rpm at room temperature (RT) for 1 h. The reaction mixture was dialyzed against 0.25 M ammonium acetate (Sigma-Aldrich) pH=5.5 containing 2 g/L chelex resin (Biorad, San Francisco, USA). The concentration of anti-MSC3-Bn-DOTA (DOTA-MSC3) was calculated from the absorption at 280 nm measured on the Infinite m200pro (Tecan, Zürich, Switzerland).

#### *Radiolabeling and Quality control*

DOTA-MSC3 was incubated with indium-111 (^111^In) chloride (Curium, Petten, the Netherlands) in 0.5 M 2-(N-morpholino)ethanesulfonic acid (MES) (Sigma-Aldrich) buffer pH 5.5 at 37°C for 30 min, with molar activities ranging from 60-180 MBq/nmol. DOTA-MSC3 was incubated with ^225^Ac chloride (van Overeem Nuclear bv, Breda, the Netherlands) in 0.1 M TRIS (Sigma-Aldrich) buffer pH 9 at 37°C for 30 min, with molar activities ranging from 45-105 kBq/nmol. Labeling efficiency and radiochemical purity were based on instant thin-layer chromatography (iTLC) on a silica gel chromatography strip (Agilent Technologies, Santa Clara, CA, USA) using 0.1 M citrate buffer (Sigma-Aldrich) pH 6 as the mobile phase. For [^111^In]In-DOTA-MSC3, iTLC was analyzed using the Typhoon FLA 7000 (GE Healthcare, Chicago, IL, USA). If labelling efficiency was <90% for *in vitro* experiments and <95% for *in vivo* experiments, the reaction mixture was purified using PD-10 desalting columns (VWR International) eluted with PBS. As ^225^Ac does not emit gamma particles it cannot be measured directly in a gamma counter. Therefore, the gamma-emitting daughter radionuclide ^221^Fr was used as a surrogate for ^225^Ac [2-4]. For [^225^Ac]Ac-DOTA-MSC3, the iTLC strip was cut in 10 pieces which were measured in a γ-counter (Wizard-2480, Perkin Elmer, Waltham, MA, USA), using the francium-221 (^221^Fr) signal as a surrogate for ^225^Ac (measured 1 h after running the iTLC, at ^221^Fr - ^225^Ac equilibrium). Radiochemical purity exceeded 95%, unless stated otherwise.

#### *Cell culture*

B16F10-OVA murine melanoma cells (kindly provided by Dr. K.L. Rock, Dana-Farber Cancer Institute, Boston [5-7]) were cultured in MEM (cat# 21575-022 Gibco, Thermo Fisher Scientific, Waltham, MA, USA) supplemented with 5% fetal calf serum (FCS), 1 mg/mL geneticin (G418), 1.5 mg/mL sodium bicarbonate, 1.5% MEM vitamins, 1% antibiotic/antimycotic, 110 µg/mL sodium pyruvate, 1% nonessential amino acids, 0.05 mM β-mercaptoethanol and 60 µg/mL hygromycin (all Gibco) at 37°C in an incubator containing 5% CO_2_. For experiments in mice, tumor cells were passaged <3 times post thawing. B16F10-OVA cells express CAIX when cultured under hypoxic conditions [8].

SK-RC-52 cells (human clear cell renal cell carcinoma, kindly provided by Memorial Sloan Kettering Cancer Center, New York, CVCL_6198) were cultured in RPMI-1640 medium (Gibco) supplemented with 2 mM glutamine (Gibco) and 10% FCS at 37°C in an incubator containing 5% CO_2_. SK-RC-52 ubiquitously overexpresses CAIX, independent of oxygen levels [9].

#### *Animal experiments*

Mice were housed in individually ventilated cages with *ad libitum* access to animal chow and water. Before starting the experiment, animals were allowed to acclimatize for at least one week. For each animal experiment, the number of animals inoculated with B16F10-OVA tumor cells was determined based on an expected tumor take of 75%. Mice with tumors outside the described tumor size range were excluded. Tumors were measured 3x/week with a caliper and tumor volumes (mm^3^) were calculated using the following formula: 4/3 * π * length/2 * height/2 *width/2. Allocation to the treatment groups was block-randomized by tumor size, using a random number generator. If possible, mice from different groups were housed together to minimize confounding effects of housing and cages were stored randomly. However, due to radiation safety reasons, mice that received ^225^Ac were housed together (although different subgroups were still randomized between the ^225^Ac cages). The order in which mice were treated, measured and examined by the biotechnicians was random, and was performed by biotechnicians blinded for group allocation. The investigator was not blinded during treatment injections, dissection and analysis.

For the therapeutic efficacy study, a priori sample size calculation was based on an effect size of 75% decline in normalized tumor volume over life-time of mice area under the curve (nAUC) compared to control, significance level of 5% and power of 80%, using one-way ANOVA. nAUC results can be found in Figure S2.

#### *Autoradiography and* *CAIX immunofluorescence*

Snap-frozen tumor sections (4 µm) were mounted on FLEX IHC Microscope Slides (Agilent Technologies). Sections were fixated with ice cold acetone at 4°C for 10 min. The slides were exposed to phospholuminescence plates in a Fujifilm BAS cassette 2025 for 1 week and scanned using Typhoon FLA 7000 bioimaging analyzer at a pixel size of 25 µm × 25 µm to obtain autoradiography images. Thereafter, slides were used for imaging of Hoechst (perfusion) and IF staining for CAIX, pimonidazole (hypoxia), 9F1 (vessels) and DAPI (nucleus), as described previously [10]. In short, before staining, slides were scanned for the fluorescent Hoechst 33342 signal. Slides were rehydrated and incubated with 5% normal donkey serum in primary antibody diluent (PAD, Biolegend, San Diego, CA, USA) at RT for 30 min and immediately incubated with 1:500 polyclonal rabbit anti-mouse CAIX (Novus Biologicals, Centennial, CO, USA) in PAD at 4°C overnight. Between consecutive steps of the staining process, sections were rinsed 3 times in PBS. Then, slides were incubated with 1:400 goat anti-rabbit FabCy3 (Jackson ImmunoResearch, Baltimore Pike, PA, USA) in PAD at 37°C for 45 min, followed by 1:400 donkey anti-goatCy3 (Jackson ImmunoResearch) in PAD at 37°C for 45 min. Sections were incubated with 9F1 (rat monoclonal antibody against mouse endothelium; Radboud University Nijmegen Medical Centre) undiluted at RT for 30 min. Next, 1:1000 rabbit anti-pimonidazole in PAD was incubated at 37°C for 30 min, followed by 1:600 donkey anti-rabbit Alexa488 (Invitrogen, Carlsbad, CA, USA) and 1:100 chicken anti-rat Alexa647 (Invitrogen) in PAD at 37°C for 30 min. Finally, sections were mounted in Fluoromount containing DAPI (Invitrogen). Tumor slides were scanned and analyzed using a digital image analysis system as described previously [11] with scanning of whole-tissue sections resulting in gray scale images.

#### *SPECT quantification*

Tumor uptake was quantified using VivoQuant Software 2021. First, SPECT scans (torso area) were loaded and resampled onto CT scans. The SPECT and CT images were manually overlayed based on the location of internal organs and the tumor. The tumor region was cropped and the volume of interest (VOI) containing the tumor was delineated based on the CT signals. Then, for each scan, a 40% threshold was determined based on the minimal and maximal SPECT value within the tumor VOI, and set as threshold to delineate the CAIX-positive (CAIX+) VOI. The CAIX-negative (CAIX-) VOI was determined by subtracting the CAIX+ VOI from the total tumor VOI. The counts in the CAIX+ and CAIX- VOIs were obtained and used to calculate the uptake in MBq/mL using a calibration factor.

#### *Calibration Factor SPECT*

The CAIX+ and CAIX- tumor volume uptake (in MBq/mL) was calculated using a calibration factor. A point source of 5 MBq ^111^In in 100 µl demi water was prepared (dose calibrator Vik202, Comecer, Joure, The Netherlands) and scanned for 60 minutes (20 bed positions, effective count time of 60 minutes, 3 minutes per bed position, spiral scan mode, normal step mode, list mode acquisition) using the GP-M 0.60 mm diameter multi-pinhole mouse collimator. The data was reconstructed using identical parameters as for the scans of the mice. The calibration factor was calculated by drawing a volume of interest in the total field of view, summing of all counts and using the following formula:

$$Calibration Factor \left( CF \right)= \frac{Activity in point source @ time of measurement \left[ MBq \right]}{\frac{\left( total counts \left[ cnts \right]*voxel size \left[ mm \right] \right)}{1000}}$$

#### *Calculation of absorbed dose values*

Dosimetry calculations for tumor and organs are based on biodistribution data (study published in [8]) and for CAIX+ and CAIX- tumor volumes are based on SPECT quantification (*in vivo* study as described in the current manuscript). These studies were performed with ^111^In-labeled MSC3. Therefore, ^111^In was used to predict the activity distribution of ^225^Ac. For ^225^Ac, a relative biological effectiveness (RBE) of 5 was used.

1) Time integrated activity coefficients (TIAC) in h/g for tumor and organs were calculated using a mono-exponential decay or association curve-fitting between the different time points in Graphpad Prism (version 5.03). If the correlation coefficient R^2^ was below 0.7, a trapezoidal approach was used [12]. Obtained TIACs were multiplied with organ S-values for ^225^Ac (OLINDA version 2.0) to calculate absorbed doses according to Medical Internal Radiation Dose (MIRD) committee methodology [13]. For tumor and lymph nodes, S-values based on GEANT4 [14] simulations of a spheric water-equivalent volume were used to calculate the absorbed dose. The S-values for an average tumor volume of 0.15 cm^3^ were obtained as previous described [15]. The absorbed dose estimations for tumor and organs with an activity dose of 15 kBq [^225^Ac]Ac-DOTA-MSC3 are shown in Table S5.

2) The TIAC (in h/mL) for CAIX+ and CAIX- tumor volumes were determined using a trapezoidal approach of the uptake values (%IA/mL) derived from the SPECT images. The obtained TIAC from each mouse was multiplied by the corresponding simulated S-values as described above to calculate absorbed doses (in Gy/kBq).

#### *Generation of dose-distribution map for ^225^Ac*

The autoradiography image of an [^111^In]In-DOTA-MSC3 injected B16F10-OVA-tumor bearing mouse was used to generated a dose-distribution map for ^225^Ac. The image data was used as a source distribution within the Monte Carlo code MCNP [16]. The decay spectra for ^225^Ac and its daughters was according to ICRP-107 [17]. The absorbed dose S-value distribution was calculated in a 50 μm voxelized structure, consisting of tissue with density of 1.02 [18]. The model was extended over 1 mm height in both Z-directions, as the autoradiography only provided data for a single central slice. The α-particle contribution to the S-values were weighted with an RBE of 5. The SPECT-based lesion TIACs were used to calculate the absorbed dose distribution according to the MIRD-schema.

### **Supplemental Tables**

Table S1: Overview of mice excluded from analysis in CD45 IF quantification.

| **Group** | **Number of mice excluded from analysis** | **Reason** |
| --- | --- | --- |
| Baseline t=0 | 2 | Tumor section too small |
| PBS 5 days p.i. | 1 | Mouse reached HEP |
| [^225^Ac]Ac-DOTA-MSC3 5 days p.i. | 2 | Tumor section too small |
| PBS 8 days p.i. | 2 | Mouse reached HEP |
| ICI 8 days p.i. | 2 | Mouse reached HEP |
| [^225^Ac]Ac-DOTA-MSC3 + ICI 8 days p.i. | 2 | Mouse reached HEP |
| **Total** | **11** |  |

Table S2: Ex vivo biodistribution analysis of [^111^In]In-DOTA-MSC3, either injected alone or in combination with [^225^Ac]Ac-DOTA-MSC3, in B16F10-OVA tumor-bearing C57Bl/6J mice (n=3/group) at 7 days p.i.. Data represent mean tissue uptake (in %IA/g ± standard deviation (SD)). A multiple unpaired t-test testing with Holm-Šídák multiple comparisons test (α = 0.05) was used to test for significant differences between [^111^In]In-DOTA-MSC3 uptake per tissue.

|  | **Tissue uptake of**  **[^111^In]In-DOTA-MSC3 (%IA/g) in mice injected with**  **[^111^In]In-DOTA-MSC3** (n=3) | **Tissue uptake of**  **[^111^In]In-DOTA-MSC3 (%IA/g) in mice injected with**  **[^111^In]In-DOTA-MSC3 and**  **[^225^Ac]Ac-DOTA-MSC3** (n=3) | **Significance** |
| --- | --- | --- | --- |
| **Blood** | 2.4 ± 0.5 | 1.3 ± 0.7 | n.s. |
| **Liver** | 17.1 ± 1.0 | 19.0 ± 0.3 | n.s. |
| **Kidneys** | 4.1 ± 0.3 | 3.3 ± 0.7 | n.s. |
| **Heart** | 1.4 ± 0.2 | 1.0 ± 0.3 | n.s. |
| **Lung** | 13.9 ± 2.8 | 9.9 ±0.9 | n.s. |
| **Thymus** | 1.8 ± 0.3 | 1.4 ± 0.3 | n.s. |
| **Brown fat** | 1.8 ± 0.2 | 1.4 ± 0.5 | n.s. |
| **Salivary gland** | 1.8 ± 0.2 | 1.4 ± 0.5 | n.s. |
| **Stomach** | 4.8 ± 1.0 | 3.5 ± 0.9 | n.s. |
| **Pancreas** | 2.8 ± 0.3 | 2.2 ± 0.5 | n.s. |
| **Spleen** | 6.3 ± 0.4 | 8.4 ± 0.3 | p = 0.0021 |
| **Duodenum** | 9.0 ± 2.8 | 6.4 ±1.1 | n.s. |
| **Jejunum** | 4.7 ± 1.1 | 5.3 ± 2.1 | n.s. |
| **Ileum** | 2.2 ± 0.4 | 2.5 ± 0.8 | n.s. |
| **Colon** | 2.1 ± 0.1 | 2.8 ± 1.2 | n.s. |
| **Lymph nodes** | 3.9 ± 0.4 | 5.2 ± 1.7 | n.s. |
| **Bone marrow** | 3.8 ± 1.0 | 3.4 ± 0.6 | n.s. |
| **Bone** | 1.7 ± 0.2 | 1.7 ± 0.6 | n.s. |
| **Muscle** | 0.3 ± 0.0 | 0.2 ± 0.0 | n.s. |
| **Tail** | 0.8 ± 0.0 | 0.8 ± 0.1 | n.s. |
| **Tumor** | 13.0 ± 4.4 | 16.1 ± 3.5 | n.s. |

Table S3: Overview of the comparisons performed in therapeutic efficacy experiment on time to reach a sixfold tumor volume relative to day 1 Kruskal-Wallis test with Dunns multiple comparisons test (α = 0.05). ICI: immune checkpoint inhibitors.

| **Comparison** | **Adjusted P-value** | **Significant** |
| --- | --- | --- |
| [^225^Ac]Ac-DOTA-MSC3 + ICI vs ICI | 0.0048 | Yes |
| non-labeled MSC3 vs PBS | >0.9999 | No |
| ICI vs PBS | >0.9999 | No |
| [^225^Ac]Ac-DOTA-MSC3 vs PBS | 0.1843 | No |
| [^225^Ac]Ac-DOTA-MSC3 + ICI vs PBS | 0.0022 | Yes |
| [^225^Ac]Ac-DOTA-MSC3 + ICI vs [^225^Ac]Ac-DOTA-MSC3 | >0.9999 | No |

Table S4: Overview of comparisons performed in therapeutic efficacy experiment on survival of Kaplan-Meier with log-rank testing and Bonferroni-corrected α of 0.00833. ICI: immune checkpoint inhibitors.

| **Comparison** | **P-value** | **Significant** |
| --- | --- | --- |
| [^225^Ac]Ac-DOTA-MSC3 + ICI vs ICI | 0.0031 | Yes |
| non-labeled MSC3 vs PBS | 0.6131 | No |
| ICI vs PBS | 0.7324 | No |
| [^225^Ac]Ac-DOTA-MSC3 vs PBS | 0.0032 | Yes |
| [^225^Ac]Ac-DOTA-MSC3 + ICI vs PBS | 0.0019 | Yes |
| [^225^Ac]Ac-DOTA-MSC3 + ICI vs [^225^Ac]Ac-DOTA-MSC3 | 0.0877 | No |

Table S5: Calculated absorbed dose (Gy) with SD for tumor and organs at risk, after injection of 15 kBq [^225^Ac]Ac-DOTA-MSC3 in B16F10-OVA tumor-bearing mice based on previously published biodistribution data [8].

|  | **Absorbed dose (Gy)** | **SD** |
| --- | --- | --- |
| **Tumor** | 81.6 | 60.2 |
| **Kidneys** | 18.5 | 6.6 |
| **Liver** | 121.0 | 31.3 |
| **Lung** | 151.5 | 33.2 |
| **Bone + bone marrow** | 2.8 | 0.3 |
| **Lymph nodes** | 17.1 | 3.3 |
| **Small intestine** | 41.5 | 13.0 |
| **Stomach** | 16.0 | 5.8 |
| **Heart** | 9.3 | 2.1 |
| **Pancreas** | 9.8 | 3.6 |
| **Spleen** | 48.8 | 11.0 |

### **Supplemental Figures**

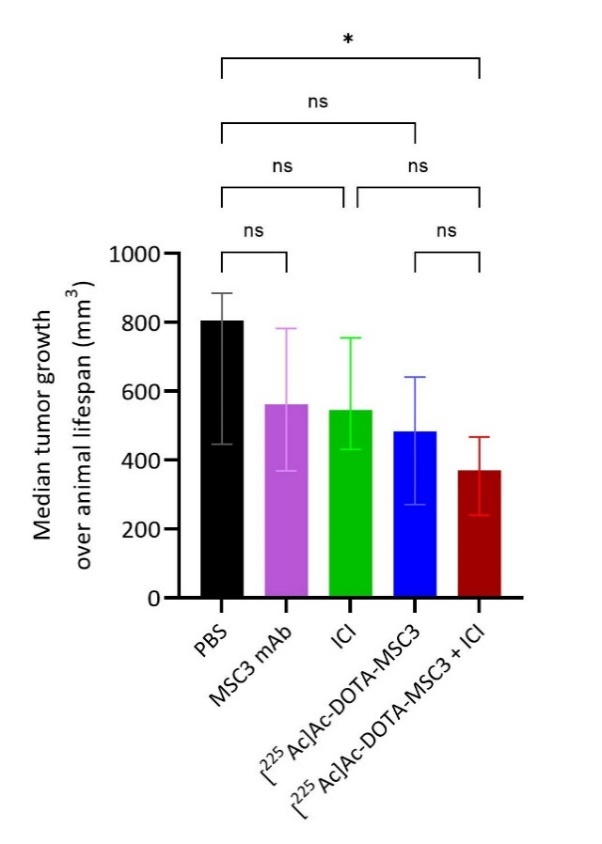

Figure S1: Median nAUC and SD with interquartile range (IQR) analyzed using Kruskal-Wallis test with Dunns multiple comparisons test.

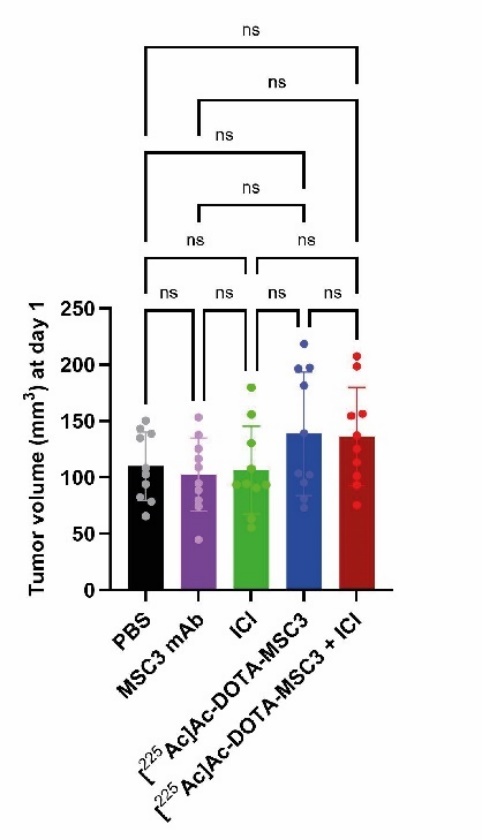

Figure S2: Mean and SD B16F10-OVA tumor sizes on day 1 after treatment with PBS control, unlabeled MSC3-DOTA, ICI, [^225^Ac]Ac-DOTA-MSC3 or [^225^Ac]Ac-DOTA-MSC3 + ICI. Differences in tumor sizes between all groups were compared using Kruskal-Wallis Test with Dunn’s multiple comparisons test, ns=not significant. ICI: immune checkpoint inhibitors.

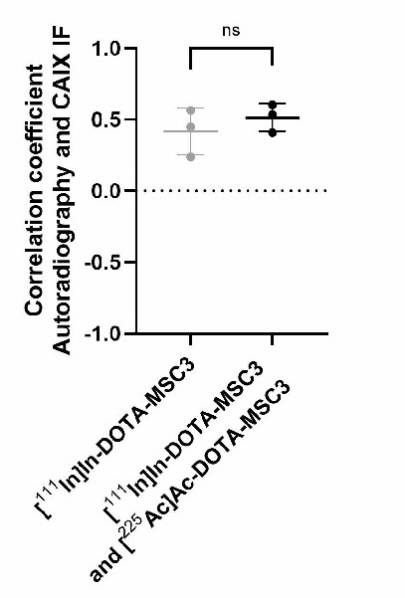

Figure S3: Spatial correlation of autoradiography and CAIX IF images expressed as correlation coefficient. Differences in correlation coefficient between B16F10-OVA tumor-bearing C57Bl/6J mice (n=3/group) injected with [^111^In]In-DOTA-MSC3 alone or in combination with [^225^Ac]Ac-DOTA-MSC3, was tested using an unpaired t-test with Welch’s correction, ns=non-significant.

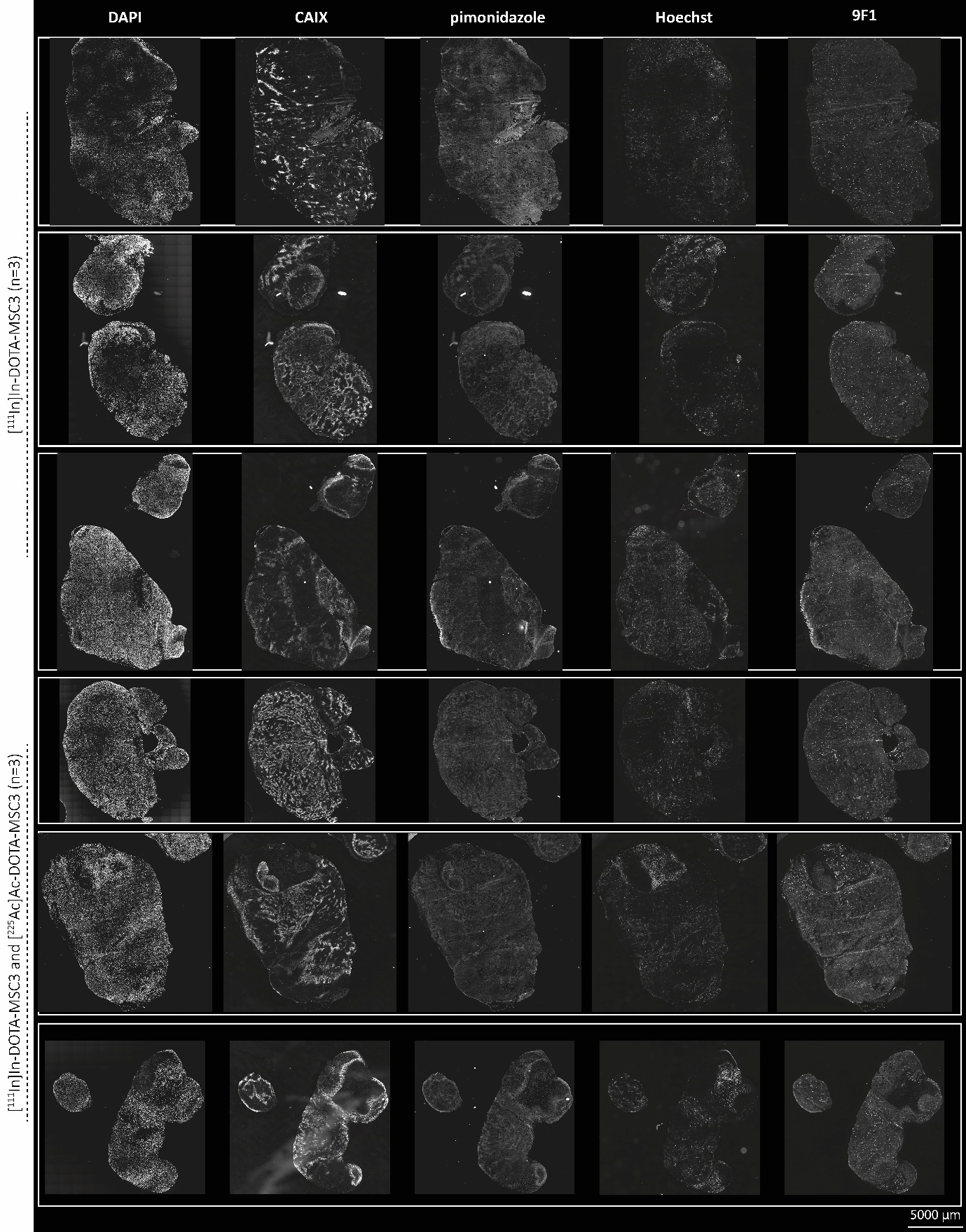

Figure S4: Overview of IF stainings in grey scale images of nuclei (DAPI), CAIX, hypoxia (pimonidazole), perfusion (Hoechst) and vessels (9F1) in tumor sections of B16F10-OVA tumor-bearing C57Bl/6J mice (n=3/group) injected with [^111^In]In-DOTA-MSC3, either injected alone or in combination with [^225^Ac]Ac-DOTA-MSC3. Brightness of gray images are adjusted for visualization purposes.

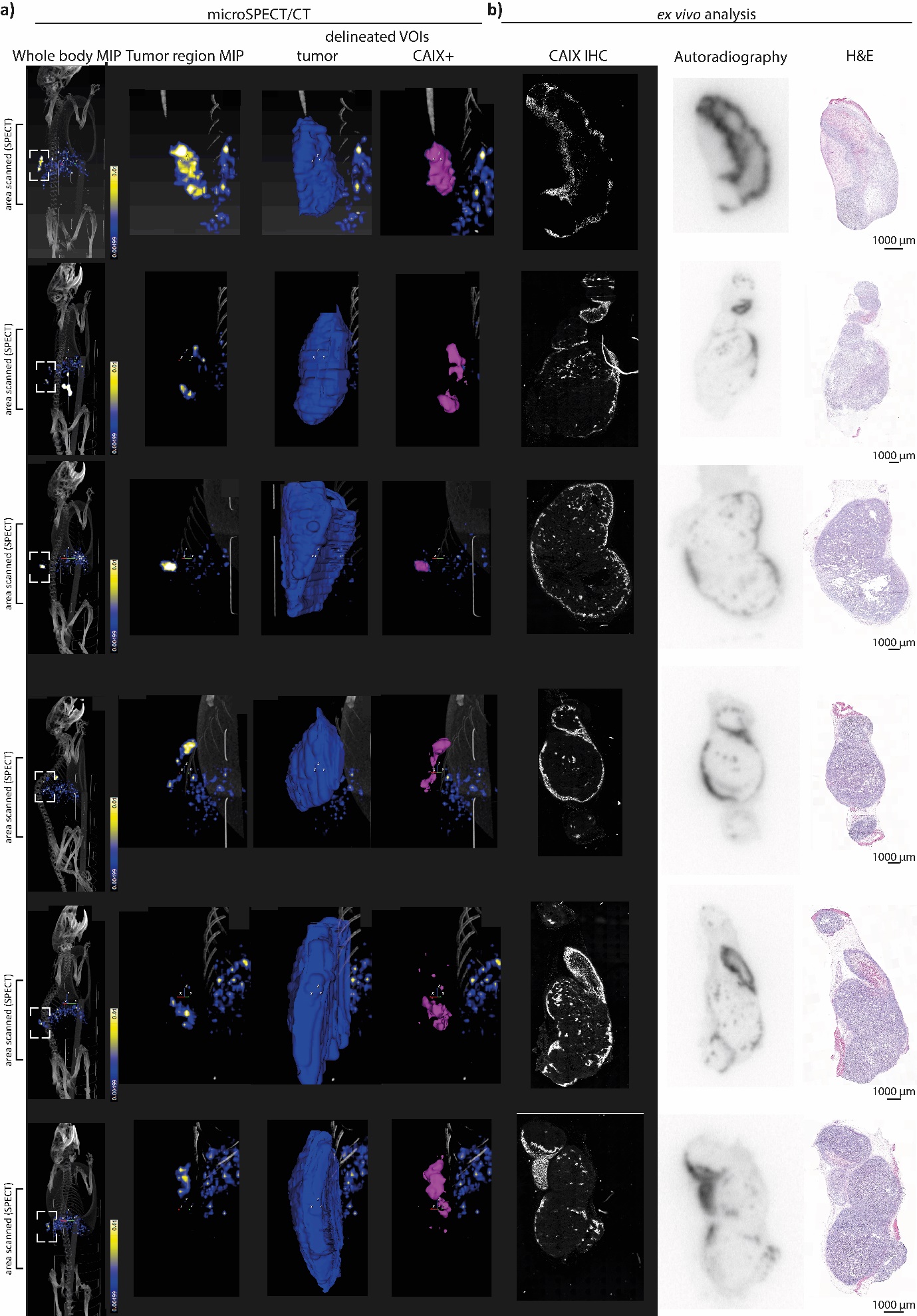

Figure S5: Accumulation of [^111^In]In-DOTA-MSC3 in B16F10-OVA tumor-bearing C57Bl/6J mice. **a)** SPECT images (torso area scanned) showing uptake at 5 days p.i.: whole body MIP, zoomed tumor region MIP, tumor area VOIs **b)** ex vivo CAIX IF, autoradiography and Hematoxylin and Eosin staining (H&E).

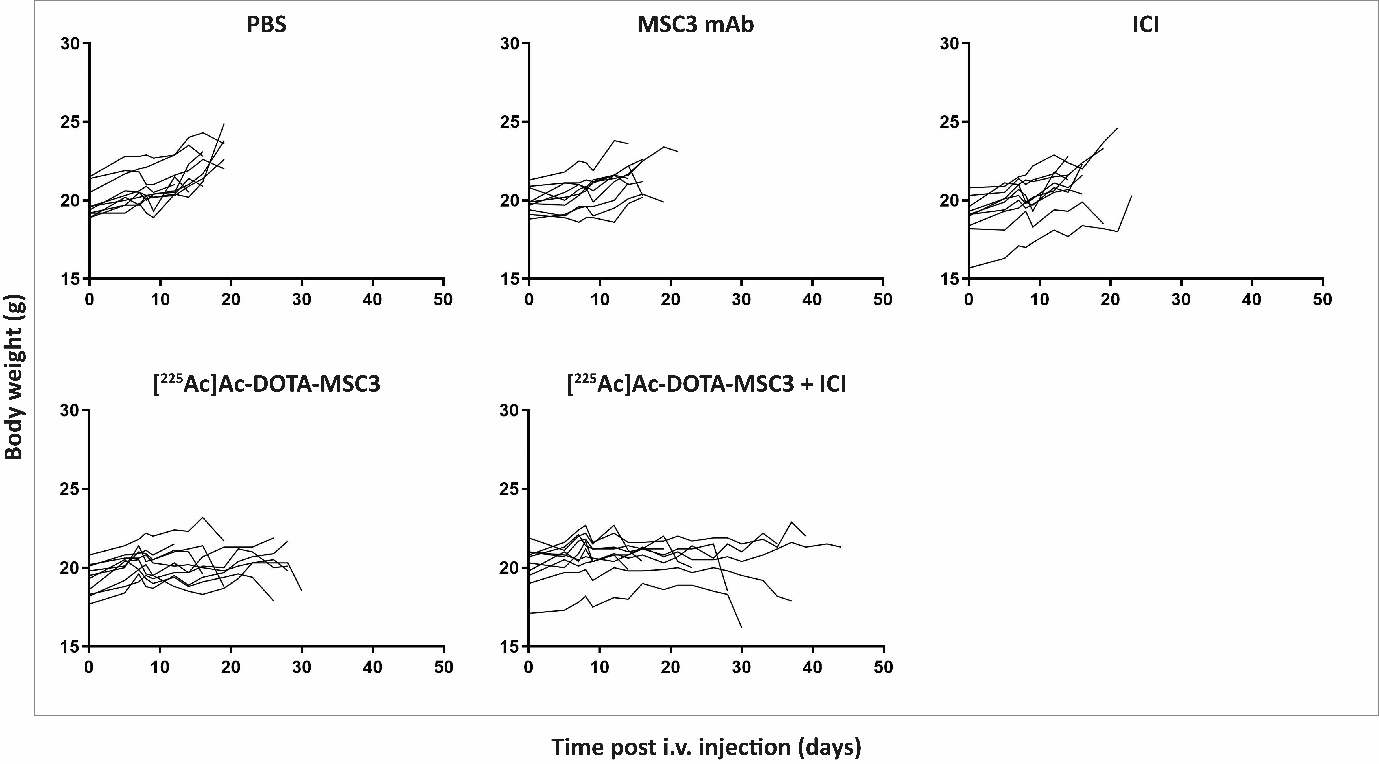

Figure S6: Body weight curves of individual B16F10-OVA tumor-bearing mice after treatment with PBS control, unlabeled MSC3-DOTA, ICI, [^225^Ac]Ac-DOTA-MSC3 or [^225^Ac]Ac-DOTA-MSC3 + ICI. ICI: immune checkpoint inhibitors.

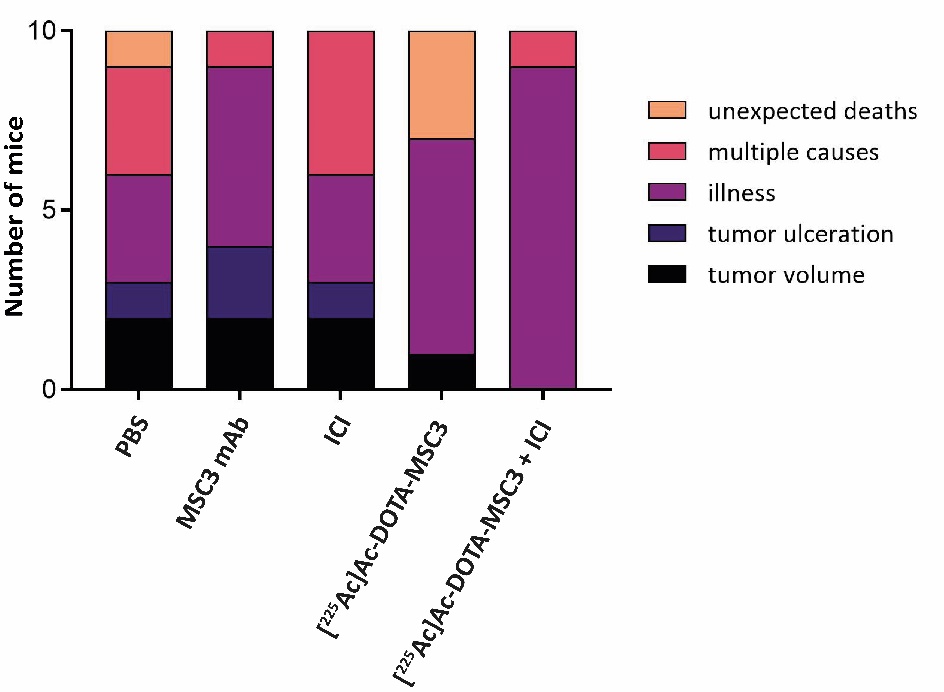

Figure S7: Overview of occurrence of humane end points in therapy experiment between the different groups. Multiple causes entails illness and tumor-related HEPs (tumor ulceration (blue) and tumor volume (black).

17.	ICRP. Nuclear decay data for dosimetric calculations. ICRP Publication 107. Ann ICRP. 2008; 38.

18.	ICRU-report-44. Tissue substitutes in radiation dosimetry and measurement; 1989.
